# Multi-modal profiling identifies CD4+CXCR5+PD-1- Tfh cells as prognostic and predictive biomarkers for response to R-CHOP therapy in human DLBCL

**DOI:** 10.1101/2024.10.10.617515

**Authors:** Sisi Yu, Hao Kong, Huaichao Luo, Xingzhong Zhao, Guanghui Zhu, Xiuqin Feng, Jie Yang, Xiangji Wu, Yu Dai, Chunwei Wu, James Q. Wang, Dan Cao, Yang Xu, Hong Jiang, Ping Wang

## Abstract

Despite the improvements in clinical outcomes for patients with Diffuse Large B-Cell Lymphoma (DLBCL), a significant proportion of those patients still face challenges with refractory/relapsed (R/R) disease after receiving first-line R-CHOP treatment. Characterizing the heterogeneity of the tumor microenvironment (TME) in diffuse large B cell lymphoma (DLBCL) is crucial for understanding relapsed/refractory disease. However, the complex and diverse nature of the TME has impeded progress. Here, using single-cell RNA sequencing(scRNA-seq), we explored the DLBCL landscape at single-cell resolution, profiling 77,344 cells from both primary and relapsed DLBCL patients. We further investigated the shared and distinct molecular and cellular features of tumor microenvironment in both primary and relapsed DLBCL tumors by integrating next-generation DNA sequencing data and multiple scRNA-seq datasets from total 72,351 cells. Our results demonstrated that there was a significant decrease of CD4+CXCR5+PD-1- Tfh cells, along with excessive activation of the TNF-NFκB signaling pathways in malignant B cells in R/R disease. Furthermore, by using multiplex IHC, we confirmed that CD4+CXCR5+PD-1- Tfh cells are prognostic and predictive biomarkers for response to R-CHOP treatment. As a proof of concept, we successfully generated mutilplexed images of CD4+CXCR5+PD-1- Tfh cells from a single DAPI staining in human DLBCL tissues through using generative artificial intelligence. Our study offers critical insights into the heterogeneity and molecular features of DLBCL, shedding light on the crucial role of CD4+CXCR5+PD-1- Tfh cells in the recurrent process.

**Highlights:**

1. Shared and distinct molecular and cellular features of tumor microenvironment in both primary and relapsed DLBCL tumors;
2. The characteristic of relapsed DLBCL TME is a dysfunctional immune microenvironment with a significant decrease of CD4+CXCR5+PD-1- Tfh cells.
3. CD4+CXCR5+PD-1- Tfh cells are prognostic and predictive biomarkers for response to R-CHOP treatment
4. Determination of CD4+CXCR5+PD-1- Tfh cells in human DLBCL tissues through using generative artificial intelligence.

## Introduction

Diffuse large B-cell lymphoma (DLBCL) is a prevalent and highly aggressive form of B cell lymphoma characterized by its high heterogeneity. Despite DLBCL being potentially curable with immunochemotherapy, such as rituximab, cyclophosphamide, doxorubicin, vincristine, and prednisone (R-CHOP), approximately 40% of patients encounter relapse and refractory^1–3^. Therefore, unraveling the underlying cause of the biological mechanism in relapsed/refractory DLBCL is a crucial clinical concern.

It is becoming increasingly recognized that acquiring a more comprehensive understanding of the tumor microenvironment (TME) in DLBCL holds promise for revealing more efficacious therapeutic approaches, even amidst the prevailing focus on investigating genetic aberrations within DLBCL tumor cells^4,5^. Multiple treatment modalities that specifically target cellular activities within the TME, including monoclonal antibodies, checkpoint blockade, and chimeric antigen receptor T cells, have either received approval or are presently under investigation for the management of aggressive B cell lymphomas^5–8^. Nevertheless, the transcriptional heterogeneity within the DLBCL tumor microenvironment (TME) that has clinical relevance is still inadequately characterized.

Recently, single-cell RNA sequencing (scRNA-seq) offers opportunities to detect and dissect the DLBCL TME^1,5^ Through this method, lymphoma-associated T cell subsets have been identified in both Follicular lymphoma and Hodgkin Lymphoma^9,10^. Few studies defined that tumor microenvironment (TME) states in DLBCL are related to drug response, chronic hepatitis B virus infection, and cell of origin (COO)^1,5,11^. However, whether a specific tumor microenvironment (TME) is associated with the relapsed/refractory DLBCL has not been clearly defined.

Here, we utilized scRNA-seq transcriptomes, DNA sequencing and multiplex IHC (mIHC) to comprehensively investigate the cellular composition of primary and relapsed/refractory DLBCL.

Through integrated analysis, we explored the shared and distinct features of relapsed/refractory DLBCL at a single-cell resolution. We identified a pivotal cellular subtype, CD4^+^ CXCR5^+^ PD-1^-^ Tfh cell, implicated in the relapse of DLBCL within the tumor microenvironment. Furthermore, we confirmed that CD4^+^ CXCR5^+^ PD-1^-^ Tfh cell is a prognostic and predictive biomarker for response to R-CHOP treatment in DLBCL. To prove a concept, we successfully generated mutilplexed images of CD4+ Tfh1 cells from a single DAPI staining in human DLBCL tissues through using generative artificial intelligence. Overall, our findings offer a comprehensive systems-level depiction of the tumor microenvironment in relapsed/refractory DLBCL.

## Methods

### Clinical information

In this study, samples for single-cell RNA sequencing and DNA sequencing were collected from 9 patients diagnosed with diffuse large B-cell lymphoma (DLBCL) who underwent surgical procedures at Sichuan Cancer Hospital in Chengdu, China. Additionally, we included a total of 32 formalin-fixed and paraffin-embedded (FFPE) DLBCL samples, obtained from patients treated with R-CHOP between January 2012 and December 2022, for mIHC analysis. Clinical and histological information was collected from the hospital’s medical record system. To ensure accuracy, two experienced pathologists independently reviewed the histology of the samples in a blinded manner and classified them based on the Ann Arbor staging system. The study protocol adhered to the principles outlined in the Declaration of Helsinki, and it was approved by the Ethical Committee of the Sichuan Cancer Hospital (SCCHEC-02-2022-009).

### Single-cell RNA sequencing

The surgical specimens were immediately collected from the operating room and kept on wet ice. Single-cell suspensions were prepared from the specimens by Tumor Dissociation Kit human (Miltenyi Biotec). The detail of the procedure is reported in the previous study^12^. Sequencing libraries were constructed using the Chromium Single Cell 3’ V3 reagent kit from 10X Genomics, following the manufacturer’s instructions. Subsequently, sequencing was carried out on the Illumina Novaseq 6000 platform, according to the manufacturer’s instructions (Illumina).

### DNA sequencing and variant calling

The details of the methods are listed in the previous study^13^. In this study, we conducted DNA sequencing utilizing previous single-cell sequencing samples. Targeted sequencing was performed on FFPE tumor samples from all patients using the NovaSeq platform (Illumina, San Diego, CA, USA). The library for sequencing was constructed using the lymphoma-associated gene mutation detection kit provided by Nanjing Shihejiyin Technology (Nanjing, China), Inc. The paired-end reads of targeted sequencing were aligned to the Human Genome Reference Consortium build 38 (GRCh38) using Burrows-Wheeler Aligner (BWA). Raw sequencing data were processed using Trimmomatic to remove adapters and low-quality reads. Tools such as Samtools, Picard, and GATK were used for BAM file handling, realignment, base recalibration, and variant calling. Mutations in the coding region were annotated using Annovar. Variants with depth >10 and variant allele frequency (VAF) >0.05 were filtered against control samples from a previous study. Non-synonymous SNVs, indels, and splicing sites were preserved based on various criteria including previous somatic mutation reports, hotspot mutations, and evidence-based categorization^13^.

### Multiplex Immunohistochemistry Staining

Multiplex IHC staining was performed using the TSA 6-color kit (M-D110061-50T, Yuanxi bio, China). 4 μm thick formalin-fixed, paraffin-embedded whole tissue sections with standard, primary antibodies sequentially and paired with TSA 6-color. Then by staining with SN470. Start the second to five rounds of staining, Slides were washed in TBST buffer and then transferred to preheated EDTA solution (100 °C) before being heat-treated using a microwave set at 20% of maximum power for 15 minutes. Slides were cooled in the same solution to room temperature. incubated with anti-CD20 (CM-0221, Jiehao bio, China) for 60 minutes and then treated with peroxidase-conjugated (HRP) secondary antibody (DS9800, Leica, Germany) for 10 minutes. Then labelling was developed for a strictly observed 10 minutes, using TSA 620 per manufacturer’s direction. Between all steps, the slides were washed with TBST buffer. The same process was repeated for the following antibodies/fluorescent dyes, in order: anti-PD-1 (10377-MM23, Sino biological, China)/ TSA 570, anti-CXCR5 (72172s, CST, US)/ TSA 520, anti-CD4 (ab133616, Abcam, US)/TSA 670, anti-CD8(BX50036, Biolynx, China)/ TSA 440. Each slide was then treated with 2 drops of SN470 (A11010-100T, Yuanxi bio, China), washed in distilled water, and manually covered slipped.

### Image analysis

The slides were visualized using the Panoramic MIDI (3DHISTECH, Hungary), and a multi-spectral image of the whole slide was scanned using a 20× objective lens. Multispectral image immixing was performed using HALO software (Indica Labs, US) image analysis with cell segmentation and spatial analysis. Briefly, in the “High-Plex FL” module, the DAPI positive cell was identified by the “cell detection” command, and each single channel intensity threshold was manually determined for each slice. We determined positive cell numbers and counted cells proportions by dividing the channel positive cell counts by all the cells detected with DAPI positive. Then all detected cells were divided into different subgroups for further analysis and folded tissue were excluded. Spatial distance is calculated by the proximity algorithm in the “Spatial analysis” module.

### Implementation of generation model

To train our model, we utilized 30 DLBCL samples, each accompanied by six IHC-stained images. Then we divided each sample into two parts: one consisting of the monochrome DAPI image, which served as the input to the model, and the other comprising the six-channel merged IHC image, which as the label for our generated outputs. Following this, we meticulously aligned the images for each sample pair and employed clustering-constrained-attention multiple-instance learning (CLAM) ^14^ method to identify cellular regions and segment patches. This process yielded 40,000 patch images, each patch with a resolution of 1024×1024 pixels. Of these, 32,000 (80%) images were designated as the training set, while the remaining 8,000 (20%) images were reserved for testing. To produce high-quality mIHC images, we employed the more robust conditional generative adversarial network (pyramid pix2pix) as our baseline model, that utlized ResNet-6Blocks as the generator(*G*) and a 5-layer PatchGAN as the discriminator(*D*)^15^. Simultaneously, we incorporated the Adaptive Supervised PatchNCE (ASP) loss function to ensure that the generated images exhibit a high degree of pixel consistency with the real images^16^.

In the training process, we deployed this model on 1 RTX 4090 GPU for training, with batchsize set to 2, and trained 50 epoches. The Adam optimizer was used with a linear decay scheduler and an initial learning rate of 2×10^-4^. The whole training process spended 4 days,

### Evaluation of generated images

In order to evaluate the performance of the model, we adopted the SSIM and PSNR to evaluate the quality of generated images. let x and y as the generated images and real image, which the SSIM and PSNRdefined as follows:

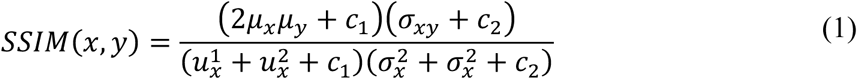

where *μ*, *σ* denoted the mean and variance of image, and *σ_xy_* is the covariance between image x and image y, *c*_1_, *c*_2_ were constant.

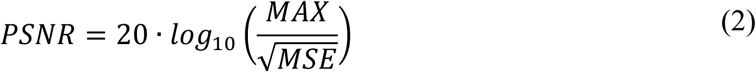

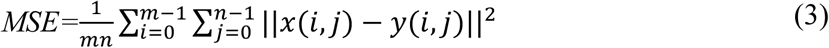

Where *MAX* was the maximum value of a pixel on an image, and m, n was the number of pixels on image x and y.

In addition, to evaluate the performance of our model in identifying CD4^+^ TFH1cells, we employed experienced doctors manually labeled 2,000 real mIHC images and generated images, and divided them into two classes: images with CD4^+^ TFH1cells and images without CD4^+^ TFH1cells. We then used metric of classificaton to assess the 2,000 generated mIHC images. First, we computed the confusion matrix, which includes the following: true positives (TP, CD4^+^ TFH1cells existed in both real and generated images), true negatives (TN, absence of CD4^+^ TFH1cells in both real and generated images), false positives (FP, CD4^+^ TFH1cells existed in generated images but not in real images), and false negatives (FN, CD4^+^ TFH1cells present in real images but missed in generated images). Then we compute the accuracy 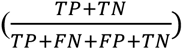, recall score 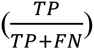, precision 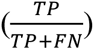, and F1 score 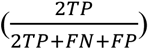

## Results

### The landscape of scRNA-seq profile of human DLBCL

To explore the cellular composition of DLBCL, we profiled clinically annotated primary or relapsed DLBCL tissues by scRNA-seq (**Fig. 1a** and **Suppl. Table 1**). After quality control (**Suppl. Table 2**), we acquired the transcriptomes of 77,344 single cells in this work. We also obtained available single-cell dataset with 72,351 cells to validate our results (**Fig. 1a**). After the first round of clustering, we found immune cells were main cell type (**Fig. S1a, b**). In combined cohort, three main cell clusters were showed in **Fig.1b**, most cells overlapped, showing the robustness of our single cell methods. In total, 21 main cell types were identified, including 3 types of B cells, 10 types of T/NK cells and 8 types of myeloid cells (**Fig. 1c**). In most cases, well-established canonical markers generally distinguished different clusters, such as *CD79A* for B cells, *CD3G* for parafollicular cells, *CD3D* for T cells, *CD68* for Myeloid cells (**Fig. S1c, d**). For cell counts of all profiles of DLBCL, top three were normal B cells (B_nor), malignant B cells (B_mal), and CD8^+^ exhausting T cells (CD8^+^_TEX) (**Fig.S1e**). The heatmap of cell type canonical genes expression and correlation of all variable gene expression matrix further confirmed the cell annotation of our work (**Fig.S1f, g)**. The immune cell UMAP figure showed the difference of cell distribution of primary and relapsed/refractory DLBCL which was located at B cell cluster (**Fig. 1d**). Bar plot illustrated the proportion of different treatment response, showing CD4^+^ Tfh, CD4^+^ TN and CD8^+^ TEMRA were enriched in primary DLBCL, and B_mal were enriched in replaced DLBCL (**Fig. 1e**). Most of enrolled subjects harbored all main cell types (**Fig. 1f)**. The ratio of observation to expectation (RO/E) displayed several specific cell types for relapse and primary DLBCL. We found that there was significant decrease of cell proportion in relapsed tumors compared to primary tumors, including CD4^+^ T cells, CD8^+^ T cells, and NK cells (**Fig. 1g**), indicating an immune dysfunctional microenvironment in relapsed tumors.

**Fig. 1.**
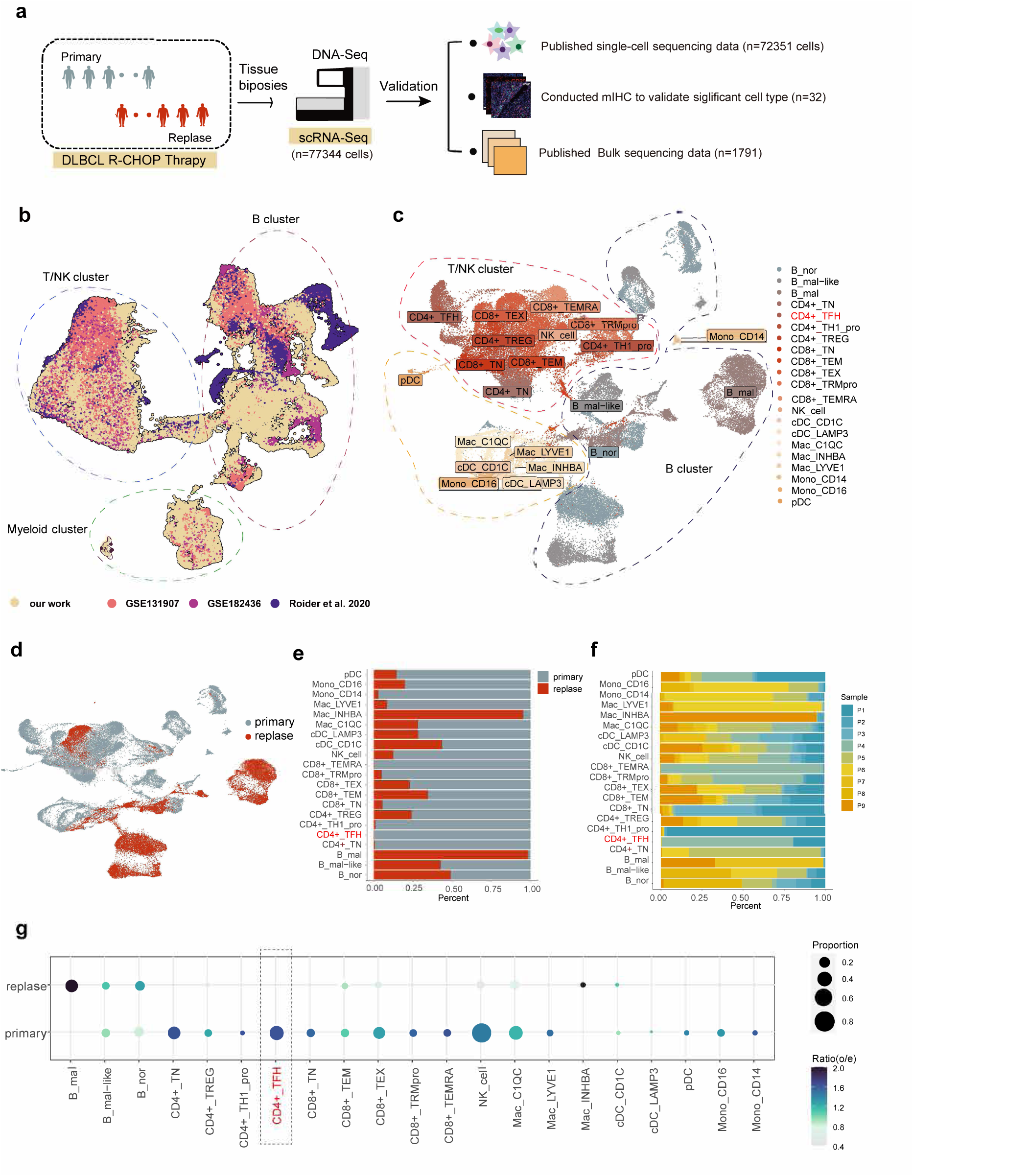
Single-cell profile of DLBCL. (a) Overview of the experiment procedure. Nine scRNA-seq data were generated from primary and replase DLBCL tissues using 10× Genomics protocol. We analyzed the transcriptome of 77344 individual cells. For significant cluster, we conducted a series of validation in published single-cell data (72352 cells), DNA-sequencing data, mIHC data (32 individuals), published bulk sequencing data (1791 individuals). (b) UMAP demonstrates the all collected DLBCL and normal lymph nodes single cell sequencing data. Dataset names were labeled on the figure. UMAP demonstrates the DLBCL single cell sequencing data in our work. Main cell types identified by graph-based clustering (c) and Treatment response (d) were labeled on the figure. Bar plot illustrating the proportion of different treatment response (e) and enrolled subjects (f) in main cell types. (g) Treatment response preference of each cell types measured by cell proportion (size) and the ratio of observed to randomly expected cell numbers (R_O/E_,color) (Zhang et al., 2018).

### Dynamic transcriptional changes for B cell cluster in human DLBCL

To further explore the dynamic transcriptional change of B cell cluster, we subseted all B cell clusters into one independent object. B cell cluster was separated into 19 clusters (**Fig. 2a**). The calculated *IGKC*/(*IGKC2*+*IGLC*) value was displayed by dot plot and UMAP (**Fig. 2b, c**). Most of the cell clusters were defined as normal or malignant B cells by previous knowledge^11^. The left clusters, which cannot be annotated, were defined as malignant-like B cells (B_mal-like). In order to validate this method, we applied the same way into combined dataset with 4 datasets (**Fig. S2a**). The B cell subgroup of this combined dataset was separated into 27 clusters (**Fig. S2b, c**). After B cell defining, all normal lymph nodes harbored normal B cells (B_nor), meanwhile all malignant B cells (B_mal) and malignant-like B cells (B_mal-like) were discovered in DLBCL group (**Fig. 2d, Fig. S2d**). By employing this method, a distinct categorization of 6 B_nor, 6 B_mal_like, and 7 B_mal clusters was achieved. (**Fig. 2e**). New B cell types percent in all B cells of one subject were displayed by boxplot, showing that B_mal cell type is the dominant cell types in relapsed group (**Fig. 2f**). B_mal and B_mal-like cell types highly expressed genes involved in the naive stage of B cell differentiation^17^ (**Fig. 2g**). Three B cell types specific genes were listed in **table S3**. Since B_mal and B_mal-like were similar, we conducted DEGs analysis between malignant group (B_mal and B_mal-like) and normal group (B_nor). These DEGs were enriched in several cancer-associated hallmark pathways (e.g., Oxidative phosphorylation, Myc targets v1, Mtorc1, TNFα signaling via NFκB (**fig. 2h**). To gain insights into the cellular progression from primary to relapsed state, we apply scTour to predict the cellular dynamics from primary to relapsed state (**Fig. 2i**)^18^. We identified three sets of differentially expressed genes (DEGs) along the primary to relapsed trajectory (**Fig. 2j**). Hypergeometric test on specific genes for each cluster along the trajectory using hyperR for C5 gene sets from MSigDB Human collections (**Fig. 2k**, and **Suppl. Table 3**). Cluster 1, consisting of leukocyte mediated immunity (*TNFRSF1B, C1QA*) decreased along the trajectory, while the other two sets, consisting of GOCC-ribosome (*RPL11, RPS8, RPL5*), increased along the trajectory. Collectively, the results demonstrated that B cells exhibited a decrease in leukocyte-mediated immunity and an increase in their proliferation ability.

**Fig. 2.**
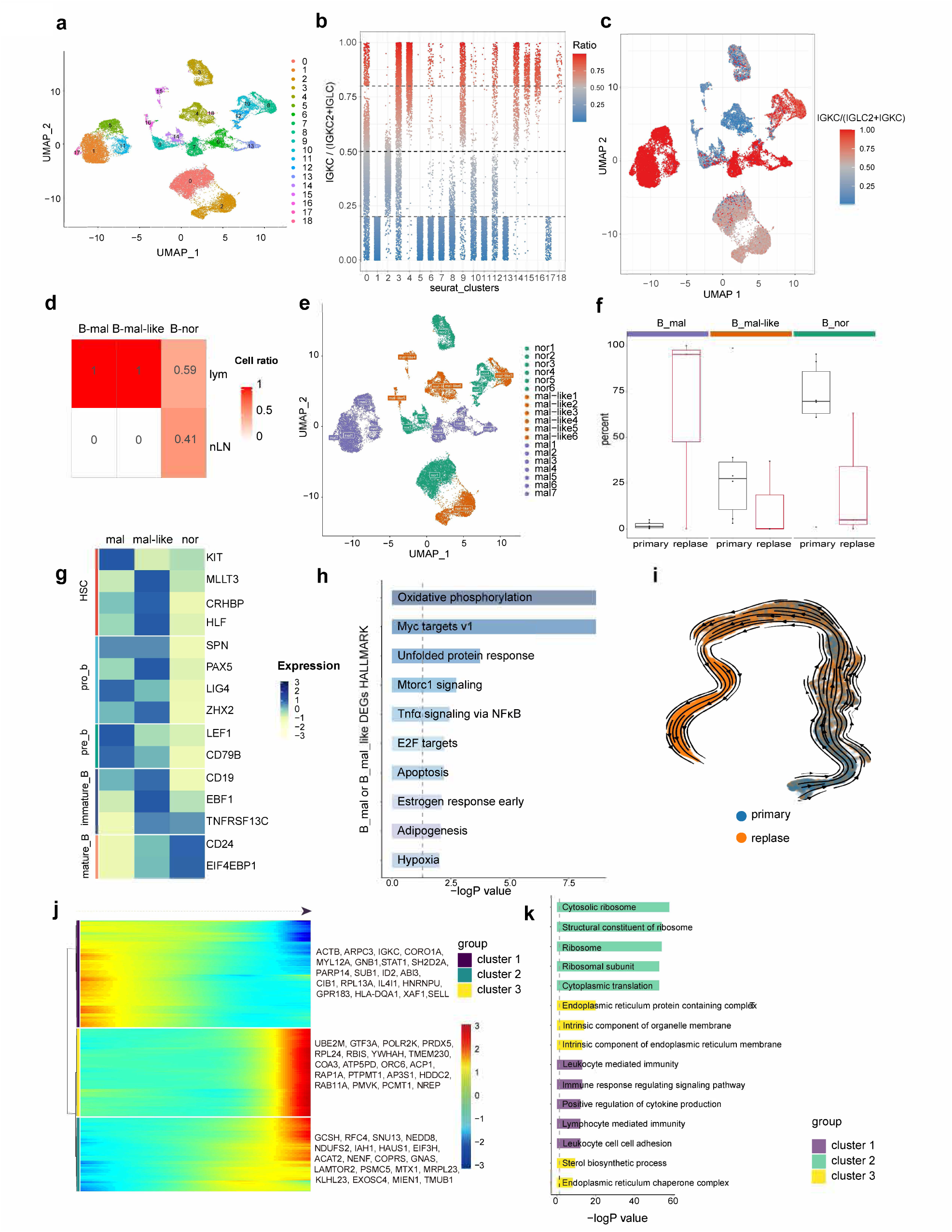
Associations of B cellular compositions with treatment group. (a) UMAP demonstrates the B cell in our DLBCL single cell sequencing data, colored by graph-based clustering. (b) We applied reported classic strategy to illustrate the strategy employed to identify malignant B cells. Each B cell was assessed for the *IGKC* fraction, calculated as *IGKC* / (*IGKC* + *IGLC2*). Most of cells with an *IGKC* fraction <0.25 or >0.75 were classified as malignant B cell, while cell clusters with various *IGKC* fraction B cells were classified as normal B cell. Unlike previous methods, we innovatively categorize clusters that fall between malignant clusters and normal cells, and have at least one quartile interval missing, as malignant-like cell clusters. *IGKC* fraction are shown by Scatter plot. (c) The scRNA expression profiles of the B cells were visualized by UMAP, colored by *IGKC* fraction. (d) To validation our method, we applied this method into combined DLBCL and normal lymph nodes datasets. We found all normal Lymph nodes B cells are classed into normal group. (e) B cell in our DLBCL single cell sequencing data were visualized using UMAP, the cells are colored by new B cell classification. (f) Relative contribution of each B cell types (%) in primary and replase biopsies. (g) Heatmap showing expression of functional genes associated with B cells differentiate in the 3 B cell cell types. (h) GSEA based on Hallmark Gene Set for DEGs comparing Malignant or Malignant like B cell versus Normal B cell. (i) scTour analysis result of B cellular dynamics from primary to replase. (j) Gene expression dynamics along the B cell trajectory. (k) hypergeometric test based on Molecular Signatures Database C5 collection for three gene cluster in B cell trajectory.

To identify the differences of genomic profile between malignant or malignant-like and normal B cells, we first inferred copy number alterations (CNAs) through the single cell–based approach. A higher burden of CNAs was identified in malignant or malignant-like B cells than in normal B cells, indicating increased chromosome instability in malignant or malignant-like B cells. Meanwhile, for relapsed DLBCL group, a higher burden of CNAs was identified in malignant than malignant-like B cells. In the 18 chromosome, malignant and malignant-like B cells shared similar CNA patterns (**Fig. S2e**). Because of the above analysis described the difference of genomic and transcript between malignant or malignant-like and normal B cells, we speculated that an upstream transcription factor (TF) might function as a key regulator of this difference. Therefore, we performed SCENIC analysis with scRNA-seq data to reconstruct co-expression modules and identify the key TF regulators for this DEGs. As a positive control, a well-established TF (i.e., *BATF*, also known as a lymphoma transcription factor) was identified as a key regulator in activated B-cell-like diffuse large B-cell lymphoma (ABC-DLBCL), which enriched in primary B_mal/B_mal-like cells (**Fig. S2f**)^19^. Other TFs (e.g., *BCL2*) were enriched in primary B_norm cells^20^. We observed that B_mal/B_mal-like cells showed enhanced activities of *NFATC1* compared to normal B cells. This TF was also enriched in relapse B_mal/B_mal-like cells, but largely absent in primary B_mal/B_mal-like and all normal B cells (**Fig. S2f**), suggesting their potential role in DLBCL relapse development. Intriguingly, *NFATC1* gene is located on chromosome 18, which is consistent with CNAs findings.

### Transcriptional fluctuation and cell fraction distribution of T cell cluster in human DLBCL

In order to validate the cell annotation process, we compared the celltypist and our method in T cell subset of combined single cell dataset, and found the result is similar (**Fig. S2g**). In combined T cell dataset, different dataset can overlap in one cell large cluster, almost all cell types in our study can be found in combined datasets (**Fig. S2h, i**).

To characterize T cell subsets in the tumor microenvironment, we have divided the T/NK large cell group, and further separated it into 19 cell types (**Fig. S3a**), the detail markers expression heatmap and UMAP labeled by the key genes are displayed in **Fig. 3b, S3a, S3b and Suppl. Table 4**. We applied T cell functional genes to explain the functional state of CD4^+^ T cells. For most cells, as T cells differentiate and mature, the naïve T cells signal decreased, and regulatory T cells (Tregs) and exhaustion T cells signal increased (**Fig. S3c**). However, it is important to note that these observations are based on specific gene expression patterns and may not necessarily reflect the overall trend at the cellular level. Furthermore, we conducted the cell fraction distribution analysis. We found that there was a significant increase of CD4^+^ Tregs in tumors from the relapsed group compared to the primary tumors, which contribute to an immunosuppressive tumor microenvironment^21^. The proportion of CD4^+^ Tfh1 (CD4^+^ CXCR5^+^ PD-1^-^) is higher in primary group than that in relapsed group (**Fig. 3c, S3d**). The distribution of CD8^+^ T cells between the two groups did not show significant differences. (**Fig. S3e)**. In the combined dataset, we found a similar pattern for CD4^+^ Tfh1 cell type (**Fig. 3d**). To further explore CD4^+^ T cells (ignored Tregs), we separated CD4^+^ T cells and found CD4^+^ Tfh1 is an independent cell cluster. DEGs (primary vs. relapse) are identified in these CD4^+^ T cells (**Fig. 3e, table S3**). The inner group stable top 10 genes are displayed in **Fig. 3f**, some published lymphoma associated genes (eg. *TCL1A* and *BCL11A*) were recognized by high expression in relapse group^22,23^. The primary DLBCL group overexpressed some innate immune associated genes (eg. *S100A8* and *PFKFB3*)^24,25^. We found the *PFKFB3* gene was highly expressed in CD4^+^ Tfh1 (**Suppl.Table 4**). Intriguingly, the canonical immune checkpoint gene *PDCD1* exhibits low and cell-specific expression in CD4^+^ Tfh1 cells (**Fig. 3g, table S4**). Hypergeometric test on the expression differences between the primary and relapse group in CD4^+^ T cells revealed that gene-expression signatures associated with innate immune system gene sets were significantly enriched in the primary DLBCL group (**Fig. 3h**). All the evidence suggests a deficiency in the innate immune microenvironment in the relapsed DLBCL group.

**Fig. 3.**
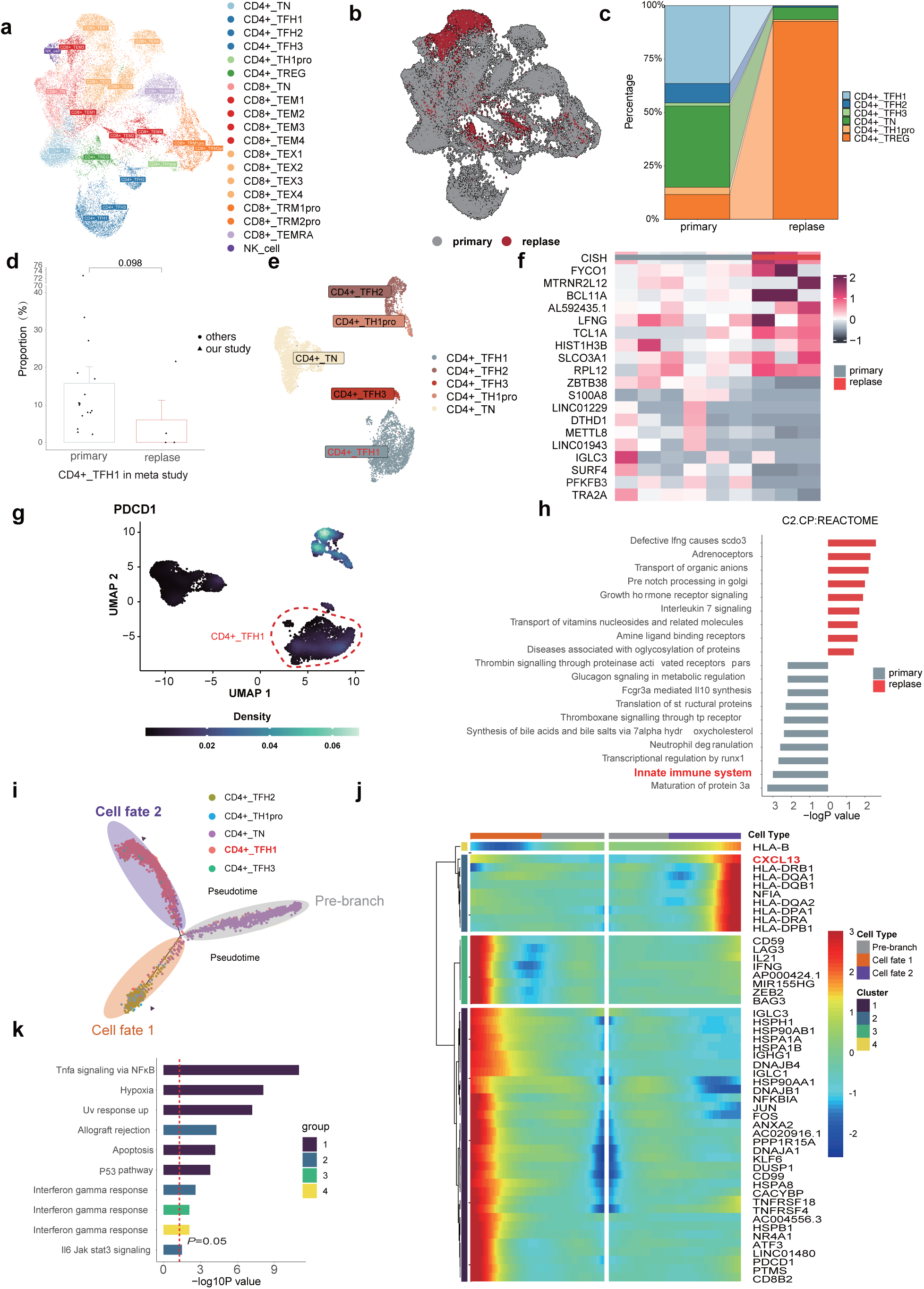
Associations of T cellular compositions with treatment group. UMAP demonstrates the T cell subset. All cell types identified by graph-based clustering (a) and Treatment response (b) were labeled on the figure. (c)Modifications in the distribution of CD4^+^T cell subset between the primary and replase groups within our dataset. (d)Percentage of CD4^+^ THF1 cell for each individual comparing primary with replase in combined DLBCL data set. (e) CD4^+^THF1 cell subset is displayed by UMAP, color by cell types. (f) Heatmap showing stable different expressed genes in CD4^+^ T cell subset between primary and replase group. (g) UMAP demonstrates CD4^+^T cell subset, colored by density of *PDCD1* gene. (h) hypergeometric test based on REACTOME pathways for stable different expressed genes. (i) Cell trajectory analysis of CD4^+^ T cell differentiation. Cells are labeled with each cell types. (j) Heatmap showing the expression changes of the 50 highly variable genes along the two branch of differentiation trajectory, color scale indicates z score. (k) Significantly enriched Hallmark annotations are shown on the bar plot.

Trajectory analysis was subsequently performed using CD4^+^ TN cells as the starting point for CD4^+^ T cell trajectory analysis with Monocle2. 2 branches and 2 cell fates were identified. In cell fate2, a higher cell density and a higher percentage of CD4^+^ Tfh1 were observed (**Fig. 3i**). By focusing on the mechanism underlying disease relapse, branch analysis was performed at branchpoint of the trajectory. A cluster 2 of genes that were highly expressed in cell fate 2 was identified, including several innate immune system related genes (*CXCL13, HLA-DB*)^26^ (**Fig. 3j**). After gene enrich analysis, the cluster 1 genes were significantly enriched on TNF-α signaling via NF-κB (**Fig. 3k**). As time progresses along the pseudo-temporal trajectory of cell fate2, the expression of TNF-α signaling via NF-κB signaling-related genes in Cluster 1 gradually decreases. The classification of myeloid cells in the single-cell profile of DLBCL is displayed in **Fig. S4**. Taken together, our data suggest that the absence of CD4^+^ Tfh1 with innate immune function in the microenvironment may be associated with relapse of DLBCL.

### Tumor cells affect the proliferation, differentiation and immune function of CD4^+^ Tfh1 cells through ligand receptor signaling and TNFα-NFκB pathway

To explore the cell communication between CD4^+^ Tfh1 cells and B cells in DLBCL, the communication analysis was performed as previously described with default database^27^. The frequencies of interactions between ligands on CD4^+^ Tfh1 cells and receptors on B_nor cells are weaker compared to those between B_mal or B_mal-like cells. Correspondingly, the frequencies of interactions between ligands on B_mal/B_mal-like cells and receptors on CD4^+^ Tfh1 cells were higher than that of B_nor cells (**Fig. S5a**). The interaction between cell membrane proteins is an important mode of communication between cells. Therefore, this study organized the or cell membrane-bound interactions on OmniPath through the liana R package, and applied these identified interaction pairs to further investigate the additional interactions between target cells. The top 20 specific membrane-bound interactions between ligands on CD4^+^ Tfh1 cells and receptors on B cells highlight the crucial role of HLA (HLA-A→APLP2; HLA-A→LILRB1, etc.) as a vital ligand in these interactions (**Fig. 4a**). APLP2 and LILRB1 may play immune inhibitory functions for innate immune responses by HLA class I molecules ^28–30^, which is consistent with previous results in Fig. 3h. The same results are replicated by CellChat tools with same database (**Fig. S5b**). Intriguingly, the top 20 specific interactions between ligands on B cells and receptors on CD4^+^ Tfh1 cells indicated that the PTPRJ-LAT pair is specific to B_mal and B_mal-like cells, while it is almost absent in the interactions between B_nor cells and CD4^+^ Tfh1 cells (**Fig. 4b**). PTPRJ-LAT may negatively regulate TCR signaling of CD4^+^ Tfh1 cells^31^.

**Fig. 4.**
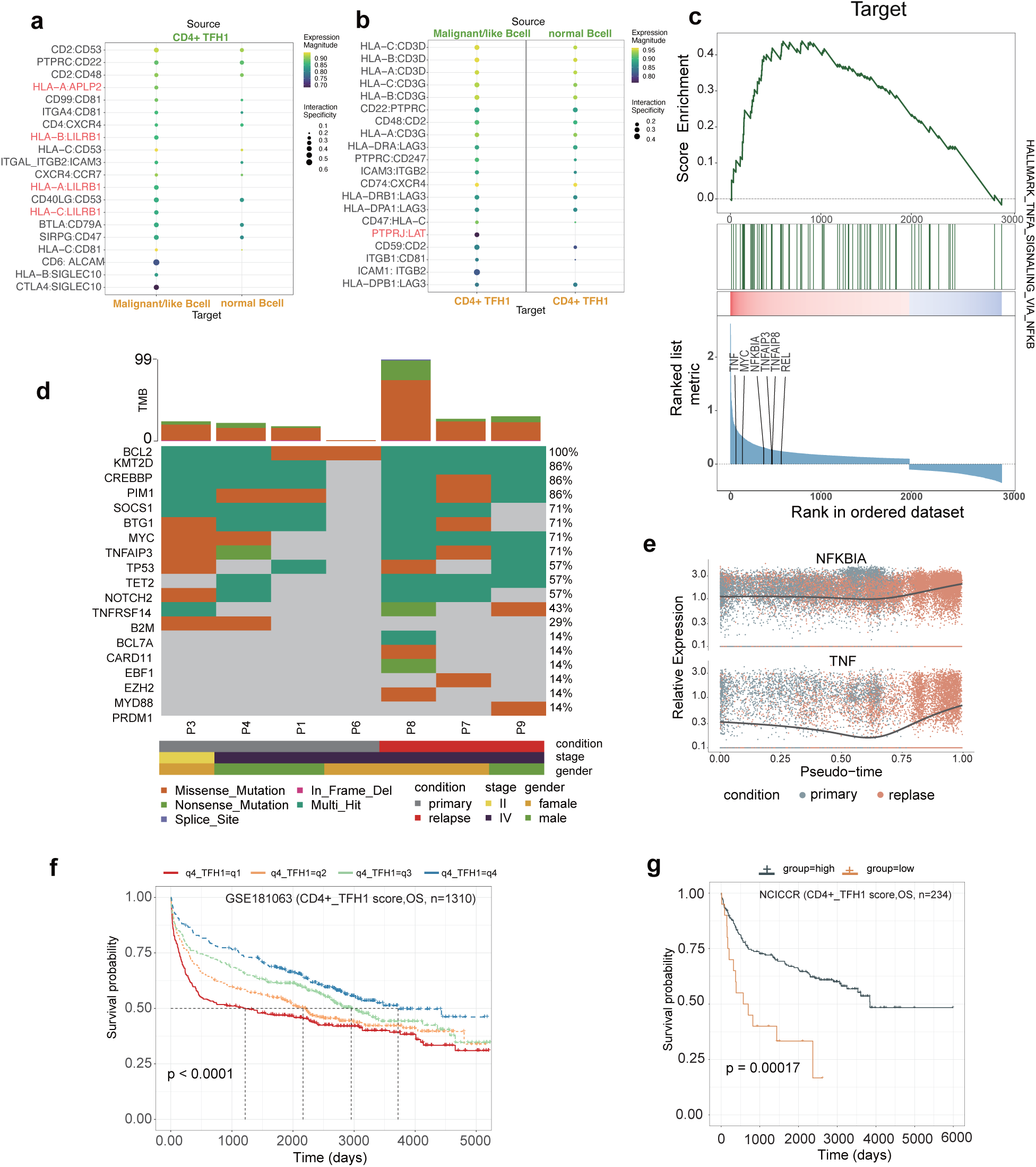
Tumor cell interact with CD4^+^TFH1 cells by TNFα-NFκB Pathway. (a, b) Dot plot showing the top 20 interaction pairs, ligands of CD4^+^TFH1 and receptors of B cell types (a), ligands of B cell types and receptors of CD4^+^TFH1(b). (c) Oncoplot of DNA sequencing showing high frequency somatic gene mutations in cosmic of DLBCL. (d) GSEA based on Malignant or Malignant like B cell upregulate genes for Hallmark TNFα signaling via NFκB gene set. (e) Two vital genes in Hallmark TNFα signaling via NFκB gene set increase as pseudotime increases. (f, g) To assess the prognostic significance of CD4^+^TFH1 in DLBCL, we conducted a study utilizing independent bulk sequence datasets. We applied Gene Set Variation Analysis (GSVA) to obtain CD4^+^TFH1-specific gene scores. Subsequently, we divided the samples into four or two groups based on quartiles (f) and median (g) and performed survival analysis using overall survival (OS, f-g).

To further study the molecular mechanisms underlying the activation of TNFα-NFκB pathway, we analyzed the genomic data of tumor samples from matched patients by DNA sequencing (**Suppl. Table 5**). The top 20 frequencies of somatic genomic signatures genes for lymphoma in COSMIC were displayed in **Fig. 4c** and **Table S4**. All patients had *BCL2* mutation, and most of patients harbored mutations in *KMI2D* and *CREBBP*. Among patients with relapsed condition, mutations in *TNFAIP3* were enriched. All the evidence suggests that the TNF-associated pathway might play an important role in the communication between CD4^+^ Tfh1 cells and neoplastic cells, as well as in the development of tumor relapse. We conducted Gene Set Enrichment Analysis (GSEA) to enrich differentially expressed genes (DEGs) between B_mal or B_mal-like cells and B_nor cells in the TNF-α signaling pathway via NF-κB. Our findings revealed that this pathway is activated in B_mal or B_mal-like cells (**Fig. 4d**). Next, we utilized the B cell trajectory analysis results from **Figure 2i** to validate the expression of two key genes (*NFKBIA* and *TNF*) in the TNF-α signaling pathway via NF-κB. We observed that the expression of these genes increased along the trajectory (**Fig. 4e**. Moreover, the crucial transcription factor *NFATC1* found in **Fig. S2f**, is associated with TNF-α Gene Expression^32,33^.

In order to validate the clinical prognostic significance of CD4^+^ Tfh1 cell types, three DLBCL bulk sequencing datasets with large sample sizes were included. Regardless of quartiles or medians, a high ssGSEA score of CD4^+^ Tfh1 cells is associated with a favorable prognosis, consistent with the findings of scRNA-seq data in this study (**Fig. 4f-g, Fig. S5c**). Taken together, the genes associated with TNF underwent alterations in both the genomic and transcriptomic levels in relapsed DLBCL tumors. Furthermore, the TNFα-NFκB signaling pathway is activated in B_mal or B_mal-like cells, potentially playing immune inhibitory functions for innate immune response of HLA on CD4^+^ Tfh1 cells, inhibiting anti-tumor immunity and promoting tumor recurrence.

### Evaluating CD4^+^ Tfh1 cell in prognosis and prediction for response to R-CHOP treatment in human DLBCL

In order to exam whether CD4^+^ Tfh1 was correlated with R-CHOP treatment response in human DLBCL, we established a multiplexed immunofluorescence panel (CD20, CD8, CD4, PD-1 and CXCR5) to characterize the interaction among neoplastic cells, T cells and CD4^+^CXCR5^+^PD-1^-^ Tfh cells in the TME (**Fig. 5a**.). 18 cases of R-CHOP responders and 14 cases of R-CHOP non-responders from DLBCL patients were enrolled for mIHC analysis as well as a clinical outcome evaluation (**table 1, Fig. S5d**). We found that the proportion of PD-1^+^ cells and CD20^+^ tumor cells were significantly higher, while there was significant decrease proportion of CD4^+^ Tfh1 in non-responders than that in responders. However, there were no changes of the proportion of total CD4^+^ T cells, CD8^+^ T cells or CXCR5^+^ cells (**Fig. 5b**). We subsequently assessed the clinical relevance of the proportion of CD4^+^CXCR5^+^PD-1^-^ Tfh cells. All patients in the entire cohort were divided into two groups based on the median proportion of CD4^+^CXCR5^+^PD-1^-^ Tfh cells. We found that DLBCL patients with a high proportion of CD4^+^CXCR5^+^PD-1^-^ Tfh cells exhibited longer overall survival (**Fig. 5c-e, Fig. S5e**). Next, we investigated the correlation of CD4^+^CXCR5^+^PD-1^-^ Tfh cells and tumor cells. There was a higher proportion of CD4^+^CXCR5^+^PD-1^-^ Tfh cells around every single CD20^+^ tumor cell in responders than in non-responders, indicating that CD4^+^CXCR5^+^PD-1^-^ Tfh cells might play a key role in the tumor progression (**Fig. S5f-g**). To better understand the spatial interaction between CD4^+^CXCR5^+^PD-1^-^ Tfh cells and tumor cells, we employed a spatial analysis by defying proximity as a percentage of CD4^+^CXCR5^+^PD-1^-^ Tfh cells which were within a radius of 50 μm from the nuclear center of CD20^+^ tumor cells (**Fig. 5f**). We found that, in R-CHOP responders, the number of CD4^+^CXCR5^+^PD-1^-^ Tfh cells within 50 μm of CD20^+^ tumor cells were significantly higher than that in non-responders. Consistently, patients with more CD4^+^CXCR5^+^PD-1^-^ Tfh cells within 50 μm of CD20^+^ tumor cells had a better survival (**Fig. 5g-h**). These results led us to examine whether CD4^+^CXCR5^+^PD-1^-^ Tfh cells could directly affect tumor cells, so we measured the number of CD4^+^CXCR5^+^PD-1^-^ Tfh cells within 20 μm to CD20^+^ tumor cells, and 20 μm represents the closest distance between two cells that have the potential to interact directly. The results showed that the number of CD4^+^CXCR5^+^PD-1^-^ Tfh cells within 20 μm of CD20^+^ tumor cells in responders were significantly more than that in non-responders (**Fig. 5i**), and the survival was significantly better among patients with more CD4^+^CXCR5^+^PD-1^-^ Tfh cells next to CD20^+^ tumor cells (**Fig. 5j-k**). All the evidence suggests that CD4^+^CXCR5^+^PD-1^-^ Tfh cells (CD4^+^ Tfh1 cells) indeed exist in DLBCL TME and could be a favorable indicator for R-CHOP treatment.

**Fig. 5.**
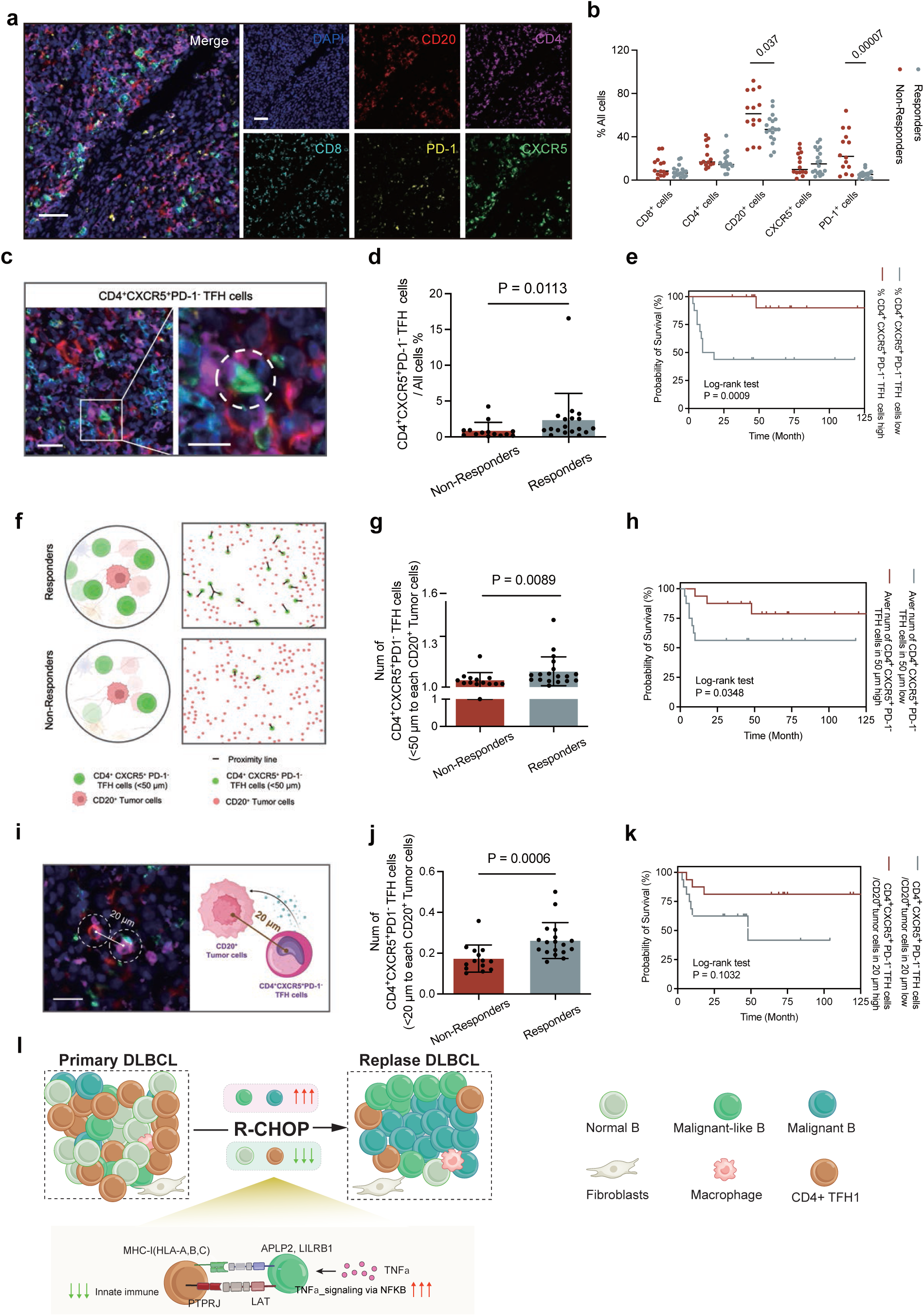
Spatial distribution of the CD4^+^CXCR5^+^PD-1^-^ TFH cells in DLBCL tumor and its correlations between patients’ survival and clinicopathological characteristics. (a) Representative composite images of DLBCL tumor tissues with multiplexed staining and imaging of individual markers (DAPI, blue; CD20, red; CD4, magenta; CD8, cyan; PD-1, yellow; CXCR5, green). Scale bar, 50 μm. (b) Percentage of PD-1^+^, CD20^+^, CD4^+^, CXCR5^+^ and CD8^+^ cells in DLBCL patients, statistical significance was determined with the multiple unpaired t test. (c) Representative images of CD4^+^CXCR5^+^PD-1^-^ TFH cells in DLBCL tumor. Scale bar, 20 μm. (d) Percentage of CD4^+^CXCR5^+^PD-1^-^ TFH cells, statistical significance was determined with the Wilcoxon signed-rank test. (e) Overall survival analysis of CD4^+^CXCR5^+^PD-1^-^ TFH cells in tissue samples from DLBCL patients. Patients were divided into two groups based on cell count above and below the median values. Differences between groups were evaluated using a log-rank test. Patients were divided into two groups based on cell count above and below the median values. Difference between groups was evaluated using a log-rank test. (f) Representative pattern (left) and representative proximity distance map (right) showing CD4^+^CXCR5^+^PD-1^-^ TFH cells within or outside the radius of 50 μm from the nuclear center of CD20^+^ tumor cells in the two groups. (g) Average number of CD4^+^CXCR5^+^PD-1^-^ TFH cells within 50 μm of CD20^+^ tumor cells in the two groups, statistical significance was determined with the Wilcoxon signed-rank test. (h) Overall survival analysis with average number of CD4^+^CXCR5^+^PD-1^-^TFH cells within 50 μm of CD20^+^ tumor cells in tissue samples from DLBCL patients. Patients were divided into two groups, based on cell count above and below the median values. Differences between groups were evaluated using a log-rank test. (i) Representative composite image of the effect of CD4^+^CXCR5^+^PD-1^-^ TFH cells on CD20^+^ tumor cells in DLBCL patients. Scale bar, 20 μm. (j) Number of CD4^+^CXCR5^+^PD-1^-^ TFH cells interacting with every single CD20^+^ tumor cells (< 20 μm) in two groups, statistical significance was determined with the Wilcoxon signed-rank test. (k) Overall survival analysis with number of CD4^+^CXCR5^+^PD-1^-^ TFH cells interacting with every single CD20^+^ tumor cells in tissue samples from DLBCL patients. Patients were divided into two groups, based on cell count above and below the median values. Differences between groups were evaluated using a log-rank test. (l) The scRNA-seq profiles reveal the heterogeneity and cellular mechanisms in relapsed/refractory DLBCL.

**Table 1.** Clinical and Pathological Characteristics of DLBCL Study Cases for Multiplexed Immunofluorescence.

| Characteristic | All patients<br>(N=32) |
| --- | --- |
| **Median age** | 54(26-81) |
| **Clinical Response** |  |
| Responders | 18 |
| Non-responders | 14 |
| **Sex** |  |
| Female | 11 |
| Male | 21 |
| **Stage** |  |
| I/II | 15 |
| III/IV | 17 |
| **IPI** |  |
| 0-2 | 23 |
| 3-5 | 9 |
| **ECOG** |  |
| 0-1 | 27 |
| 2-3 | 5 |
| **Molecular subtype** |  |
| Non-GCB | 22 |
| GCB | 10 |
| **Elevated LDH** |  |
| Low | 16 |
| High | 16 |
| **B-symptoms** |  |
| Presence | 7 |
| Absence | 25 |
| **Extranodal site** |  |
| 0-1 | 21 |
| >1 | 11 |

### Prediction and identification of CD4^+^ TFH1 cell using a deep generative network model

We have identified CD4^+^ TFH1 cells as a predictive biomarker for R-CHOP treatment response in human DLBCL. However, the detection of CD4^+^ TFH1 cells relied on multiplex immunohistochemistry (mIHC) images, which were challenging to obtain under standard conditions. To streamline the acquisition of mIHC images, we developed an end to end conditional generative adversarial network (GAN) for predicting mIHC images from DAPI-stained images. The model was trained on approximately 32,000 paired patch images (**Methods, Fig.6a**) and exhibited robust performance, achieving a structural similarity index measure (SSIM) of 0.69 and a peak signal-to-noise ratio (PSNR) of 24.83 in the testing dataset (**Methods**, N=8,000).

To further evaluate the model’s capability in detecting CD4^+^ TFH1 cells, we manually labeled 2,000 real mIHC images from the testing dataset, of which 420 contained CD4^+^ TFH1 cells (Methods). We then used these labels (images with CD4^+^ TFH1 cells were considered positive samples) to evaluate the generated mIHC images. The results showed that the model demonstrated an accuracy of 0.86 and a recall score of 0.68 in recognizing CD4^+^ TFH1 cells (**Methods, Fig.6b**). As shown in **Figure 6c**, the generated mIHC images effectively detected CD4^+^ TFH1 cells in the corresponding sample positions. This indicates that our model can efficiently generate mIHC images capable of accurately identifying CD4^+^ TFH1 cells.

**Fig. 6.**
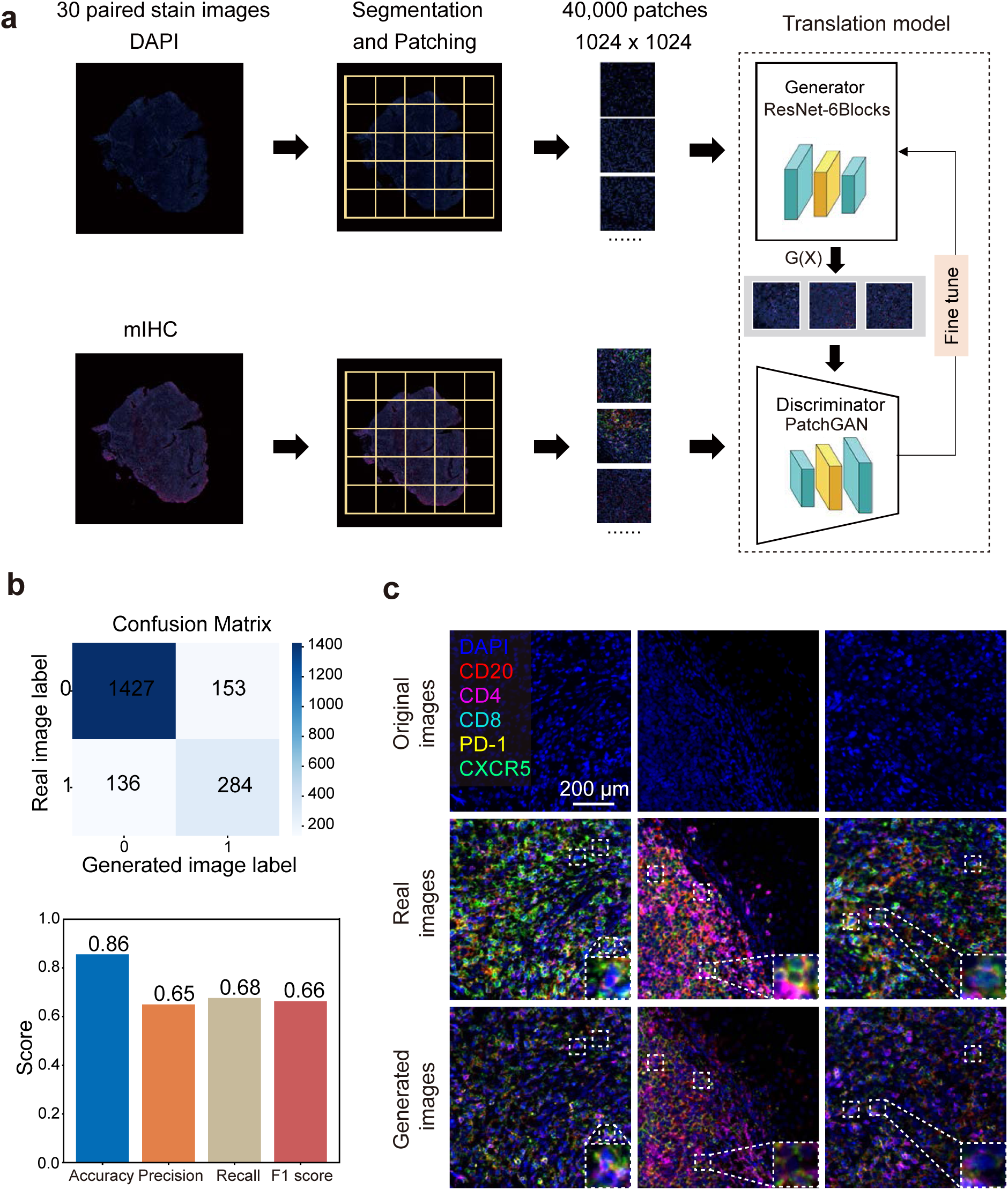
Prediction of mIHC Images from DAPI-Stained Images and Subsequent CD4+ TFH1 Cell Classification. (a) Schematic diagram of the mIHC image prediction network. The generation network comprises a generator and a discriminator. The generator utilizes a ResNet-6block architecture to produce mIHC images, while the discriminator employs a PatchGAN-based modalities discriminator module. (b) Classification performance for CD4^+^ TFH1 cells using generated mIHC images. The upper panel displays the confusion matrix for CD4^+^ TFH1 cell classification across 2000 generated mIHC images. The lower panel presents various classification metrics for CD4^+^ TFH1 cell identification, including accuracy 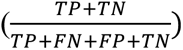, recall score 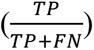, precision 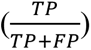, and F1 score 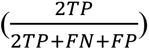. (c) The visualization of image generation results. Here we show three cases. In each column, the top image is the DAPI-stained Image, the middle image is the corresponding mIHC image, and the bottom one is the synthetic mIHC image generated with our proposed method (DAPI, blue; CD20, red; CD4, magenta; CD8, cyan; PD-1, yellow; CXCR5, green), Scale bar, 200 μm.

## Discussion

In this study, we provided a comprehensive understanding of the cellular landscape of DLBCL, spanning from the primary diagnosis to the relapsed state following R-CHOP treatment. By utilizing scRNA-seq transcriptomes, next-generation DNA sequencing, Multiplex IHC and multiple DLBCL datasets, we successfully validated our findings. This approach allowed us to depict the characteristics of tumor cells and their associated immune microenvironments, shedding light on the changes occurring in cellular and molecular networks throughout the disease progression and treatment response of DLBCL.

The importance of the tumor microenvironment (TME) in DLBCL has been increasingly recognized^34–36^. The composition of the tumor microenvironment (TME) and its interaction with malignant cells can potentially provide insights into the significance of various genes, such as CXCL13^37^, PD-1^36^, DLBCL subtype-associated genes (such as *TNFAIP3*, *CARD11*, *CD79B*, and *REL*)^36^, as well as genes associated with Tfh cells (*CD4, CXCR5*, and *ICOS*)^38^, in terms of immune evasion and the promotion of tumor progression. The TME of DLBCL is a complex entity, encompassing a variety of components, such as T and B lymphocytes, tumor-associated macrophages (TAMs), myeloid-derived suppressor cells (MDSCs), cancer-associated fibroblasts (CAFs), tumor-associated neutrophils (TANs), NK cells and other cellular elements^5,38,39^. In our findings, we have identified T and B lymphocytes as crucial cell types in the context of relapsed/refractory DLBCL with R-CHOP treatment. Specifically, CD4+ Tfh1 cell could be a predictive biomarker for R-CHOP treatment response in human DLBCL.

In this study, we identified a novel subtype of Tfh cells (CD4^+^ Tfh1, CD4^+^ CXCR5^+^ PD-1^-^ Tfh), which exhibited low levels of infiltration in the relapsed group and was associated with a favorable prognosis in DLBCL. Similar to Tfh X13 and GC Tfh cells, this cell type also produces significant amounts of CXCL13, which is a major ligand involved in interactions with B_mal or B_mal-like cells. From naive T cells, one specific cell fate is the differentiation into CD4^+^ Tfh1 cells, which exhibit high innate immune function. In R-CHOP-treated DLBCL, the presence of "cold" tumor features characterized by a low proportion of T cells has been associated with an unfavorable prognosis^34^. We believe that in DLBCL with unfavorable prognosis, the absence of CD4^+^ Tfh1 (CD4^+^ CXCR5^+^ PD-1^-^ Tfh, **Fig. 5l**), plays a crucial role in disease relapse.

Through systemic multi-patient analyses, we have uncovered malignant molecular features of DLBCL. Our findings elucidate the existence of three major B cell types in DLBCL (B_mal, B_mal-like, B_nor). As B cells progress from B_nor to B_ mal-like to B_mal, they undergo a dedifferentiation process, and the expression of genes associated with hematopoietic stem cells gradually increases (**Fig. 5l**). The proportion of B_mal was specifically associated with relapse condition. ABC DLBCL subtype-associated genes mainly involved NF-kB pathway ^36^, we observed NF-κB overactivation in tumor cells (B_mal and B_mal-like). It is possible that malignant B cell TNF act as a ligand and interacts with the ICOS/ICOSL ligand-mediated interaction in CD4^+^ Tfh cells. We also identified pronounced and significant copy number variations on chromosome 18 in relapse group B_mal and B_mal-like cells. Furthermore, we observed enhanced activity of the transcription factor NFATC1, located on chromosome 18, in both of these cell types. Additionally, NFATC1 has been identified as a critical signaling gene in the NF-κB pathway^40^. Further investigations will be needed to uncover the underlying mechanisms. Collectively, our findings indicate an association between the B-cell type involved in relapse and specific chromosomal abnormalities as well as transcription factor alterations. These observations may provide partial insight into the mechanisms underlying refractory and recurrence DLBCL following R-CHOP treatment.

Our study developed a deep generative network model to predicting mIHC images from DAPI-stained images. This method addressed the challenge of detecting CD4+ TFH1 cells, a predictive biomarker for R-CHOP treatment response in human DLBCL, without the need for complex mIHC procedures. Our generative model adopted ASP loss to keep the consistencies between the generated images and real images at pixel level compared with common conditional GAN model. The model’s efficacy in detecting CD4+ TFH1 cells was validated with 2,000 manually labeled mIHC images, achieving an accuracy of 0.86 and a recall score of 0.68. These results underscore the model’s potential in accurately identifying CD4+ TFH1 cells, as evidenced by the effective detection in corresponding sample positions. However, our model focused on generating mIHC images rather than directly classifying CD4^+^ TFH1 cell types, while the accuracy of CD4^+^ TFH1 cell identification requires further improvement by training a novel detection model based on cell type annotation data. In conclusion, our generative model offered a promising solution for generating high-quality mIHC images from DAPI-stained inputs, facilitating the identification of CD4+ TFH1 cells and potentially improving clinical outcomes in DLBCL treatment.

Our study has several notable limitations. Firstly, the sample size was limited, which may restrict the generalizability of our findings. Secondly, the intricate interplay between malignant B cell TNF and ICOS/ICOSL signaling in CD4+ Tfh cells requires further validation through rigorous animal and cell experiments to elucidate the underlying mechanisms. Despite some limitations, this study represents one of the largest single-cell studies conducted on both primary and relapsed DLBCL. Additionally, we utilized nearly all available scRNA-seq data to date and some large-scale bulk sequencing datasets of DLBCL to validate our main findings. Moreover, we validated our findings through immunofluorescence staining on DLBCL tissue samples, including those from both responders and non-responders following R-CHOP therapy.

In summary, we characterized the shared and distinct features of primary and relapsed DLBCL patients at single-cell resolution. The deficiency of CD4^+^ CXCR5^+^ PD-1^-^ Tfh cells with innate immune functionality and the activation of TNF-NFκB signaling in malignant B cells are vital underlying cellular-level mechanisms for the tumor microenvironment in relapsed/refractory DLBCL.

## Declaration of Human Participant

The study protocol adhered to the principles outlined in the Declaration of Helsinki, all written informed consent from the participants/patients in this study were obtained and it was approved by the Ethical Committee of the Sichuan Cancer Hospital (SCCHEC-02-2022-009).

## Data Availability Declaration

The raw ScRNA-SEQ data supporting this study have been deposited at National Genomics Data Center (NGDC, https://ngdc.cncb.ac.cn/) at PRJCA026384. Access to the data is subject to approval and a data sharing agreement due to ethical and legal restrictions.

## Code availability

Codes and pretrained model for this work are available via GitHub at https://github.com/ScuJianglab/Jianglab_DLBCL/.

## Acknowledgements

We would like to thank all members of the Hong Jiang laboratory (Sichuan University) for insightful comments regarding this work. We would like to thank Dr. Yu Liu (Sichuan University) for proofreading the manuscript and helpful discussion. We acknowledge the Core Facilities of Sichuan University.

## Author Contributions

PW, HJ and YX designed the research; DC supervised clinical studies; SSY, PW and HJ wrote the paper; SSY and DC collected clinical samples and analyzed data; HCL and HJ analyzed scRNA data; SSY and HK performed mIHC and analyzed data; JY, XJW, YD and CWW analyzed data. XZ and GZ developed the generative adversarial model and predicted mIHC images. JQW proofread the paper.

## Reference

1 Ye, X. et al. A single-cell atlas of diffuse large B cell lymphoma. Cell Rep 39, 110713 (2022). 10.1016/j.celrep.2022.110713

2 de Charette, M. & Houot, R. Hide or defend, the two strategies of lymphoma immune evasion: potential implications for immunotherapy. Haematologica 103, 1256–1268 (2018). 10.3324/haematol.2017.184192

3 Tilly, H. et al. Diffuse large B-cell lymphoma (DLBCL): ESMO Clinical Practice Guidelines for diagnosis, treatment and follow-up. Ann Oncol 26 **Suppl 5**, v116–125 (2015). 10.1093/annonc/mdv304

4 Scott, D. W. & Gascoyne, R. D. The tumour microenvironment in B cell lymphomas. Nat Rev Cancer 14, 517–534 (2014). 10.1038/nrc3774

5 Steen, C. B. et al. The landscape of tumor cell states and ecosystems in diffuse large B cell lymphoma. Cancer Cell 39, 1422–1437.e1410 (2021). 10.1016/j.ccell.2021.08.011

6 Advani, R. et al. CD47 Blockade by Hu5F9-G4 and Rituximab in Non-Hodgkin’s Lymphoma. N Engl J Med 379, 1711–1721 (2018). 10.1056/NEJMoa1807315

7 Armand, P. et al. Programmed Death-1 Blockade With Pembrolizumab in Patients With Classical Hodgkin Lymphoma After Brentuximab Vedotin Failure. J Clin Oncol 34, 3733–3739 (2016). 10.1200/jco.2016.67.3467

8 Schuster, S. J. et al. Tisagenlecleucel in Adult Relapsed or Refractory Diffuse Large B-Cell Lymphoma. N Engl J Med 380, 45–56 (2019). 10.1056/NEJMoa1804980

9 Aoki, T. et al. Single-Cell Transcriptome Analysis Reveals Disease-Defining T-cell Subsets in the Tumor Microenvironment of Classic Hodgkin Lymphoma. Cancer Discov 10, 406–421 (2020). 10.1158/2159-8290.Cd-19-0680

10 Andor, N. et al. Single-cell RNA-Seq of follicular lymphoma reveals malignant B-cell types and coexpression of T-cell immune checkpoints. Blood 133, 1119–1129 (2019). 10.1182/blood-2018-08-862292

11 Roider, T. et al. Dissecting intratumour heterogeneity of nodal B-cell lymphomas at the transcriptional, genetic and drug-response levels. Nat Cell Biol 22, 896–906 (2020). 10.1038/s41556-020-0532-x

12 Corgnac, S., Lecluse, Y. & Mami-Chouaib, F. Isolation of tumor-resident CD8(+) T cells from human lung tumors. STAR Protoc 2, 100267 (2021). 10.1016/j.xpro.2020.100267

13 Zhang, M. C. et al. Genetic subtype-guided immunochemotherapy in diffuse large B cell lymphoma: The randomized GUIDANCE-01 trial. Cancer Cell 41, 1705–1716.e1705 (2023). 10.1016/j.ccell.2023.09.004

14 Lu, M. Y. et al. Data-efficient and weakly supervised computational pathology on whole-slide images. Nat Biomed Eng 5, 555–570 (2021). 10.1038/s41551-020-00682-w

15 Liu, S. et al. in 2022 IEEE/CVF Conference on Computer Vision and Pattern Recognition Workshops (CVPRW). 1814–1823.

16 Li, F., Hu, Z., Chen, W. & Kak, A. in International Conference on Medical Image Computing and Computer-Assisted Intervention. 632–641 (Springer).

17 Morgan, D. & Tergaonkar, V. Unraveling B cell trajectories at single cell resolution. Trends Immunol 43, 210–229 (2022). 10.1016/j.it.2022.01.003

18 Li, Q. scTour: a deep learning architecture for robust inference and accurate prediction of cellular dynamics. Genome Biol 24, 149 (2023). 10.1186/s13059-023-02988-9

19 Care, M. A. et al. SPIB and BATF provide alternate determinants of IRF4 occupancy in diffuse large B-cell lymphoma linked to disease heterogeneity. Nucleic Acids Res 42, 7591–7610 (2014). 10.1093/nar/gku451

20 Ci, W., Polo, J. M. & Melnick, A. B-cell lymphoma 6 and the molecular pathogenesis of diffuse large B-cell lymphoma. Curr Opin Hematol 15, 381–390 (2008). 10.1097/MOH.0b013e328302c7df

21 Spasevska, I. et al. Diversity of intratumoral regulatory T cells in B-cell non-Hodgkin lymphoma. Blood Adv 7, 7216–7230 (2023). 10.1182/bloodadvances.2023010158

22 Aggarwal, M. et al. TCL1A expression delineates biological and clinical variability in B-cell lymphoma. Mod Pathol 22, 206–215 (2009). 10.1038/modpathol.2008.148

23 Beà, S. et al. Clinicopathologic significance and prognostic value of chromosomal imbalances in diffuse large B-cell lymphomas. J Clin Oncol 22, 3498–3506 (2004). 10.1200/jco.2004.11.025

24 Pham, D. L. et al. Serum S100A8 and S100A9 Enhance Innate Immune Responses in the Pathogenesis of Baker’s Asthma. Int Arch Allergy Immunol 168, 138–146 (2015). 10.1159/000441678

25 Jiang, H. et al. PFKFB3-Driven Macrophage Glycolytic Metabolism Is a Crucial Component of Innate Antiviral Defense. J Immunol 197, 2880–2890 (2016). 10.4049/jimmunol.1600474

26 Crux, N. B. & Elahi, S. Human Leukocyte Antigen (HLA) and Immune Regulation: How Do Classical and Non-Classical HLA Alleles Modulate Immune Response to Human Immunodeficiency Virus and Hepatitis C Virus Infections? Front Immunol 8, 832 (2017). 10.3389/fimmu.2017.00832

27 Dimitrov, D. et al. Comparison of methods and resources for cell-cell communication inference from single-cell RNA-Seq data. Nat Commun 13, 3224 (2022). 10.1038/s41467-022-30755-0

28 Baía, D. et al. Interaction of the LILRB1 inhibitory receptor with HLA class Ia dimers. Eur J Immunol 46, 1681–1690 (2016). 10.1002/eji.201546149

29 Tuli, A. et al. Amyloid precursor-like protein 2 association with HLA class I molecules. Cancer Immunol Immunother 58, 1419–1431 (2009). 10.1007/s00262-009-0657-z

30 Zeller, T. et al. Perspectives of targeting LILRB1 in innate and adaptive immune checkpoint therapy of cancer. Front Immunol 14, 1240275 (2023). 10.3389/fimmu.2023.1240275

31 Baker, J. E., Majeti, R., Tangye, S. G. & Weiss, A. Protein tyrosine phosphatase CD148-mediated inhibition of T-cell receptor signal transduction is associated with reduced LAT and phospholipase Cgamma1 phosphorylation. Mol Cell Biol 21, 2393–2403 (2001). 10.1128/mcb.21.7.2393-2403.2001

32 Kaminuma, O. et al. Differential contribution of NFATc2 and NFATc1 to TNF-alpha gene expression in T cells. J Immunol 180, 319–326 (2008). 10.4049/jimmunol.180.1.319

33 Yarilina, A., Xu, K., Chen, J. & Ivashkiv, L. B. TNF activates calcium-nuclear factor of activated T cells (NFAT)c1 signaling pathways in human macrophages. Proc Natl Acad Sci U S A 108, 1573–1578 (2011). 10.1073/pnas.1010030108

34 Song, J. Y. et al. Low T-cell proportion in the tumor microenvironment is associated with immune escape and poor survival in diffuse large B-cell lymphoma. Haematologica 108, 2167–2177 (2023). 10.3324/haematol.2022.282265

35 Merdan, S. et al. Gene expression profiling-based risk prediction and profiles of immune infiltration in diffuse large B-cell lymphoma. Blood Cancer J 11, 2 (2021). 10.1038/s41408-020-00404-0

36 Cioroianu, A. I. et al. Tumor Microenvironment in Diffuse Large B-Cell Lymphoma: Role and Prognosis. Anal Cell Pathol (Amst) 2019, 8586354 (2019). 10.1155/2019/8586354

37 Liu, B., Zhang, Y., Wang, D., Hu, X. & Zhang, Z. Single-cell meta-analyses reveal responses of tumor-reactive CXCL13(+) T cells to immune-checkpoint blockade. Nat Cancer 3, 1123–1136 (2022). 10.1038/s43018-022-00433-7

38 Ng, W. L., Ansell, S. M. & Mondello, P. Insights into the tumor microenvironment of B cell lymphoma. J Exp Clin Cancer Res 41, 362 (2022). 10.1186/s13046-022-02579-9

39 Liu, Y., Zhou, X. & Wang, X. Targeting the tumor microenvironment in B-cell lymphoma: challenges and opportunities. J Hematol Oncol 14, 125 (2021). 10.1186/s13045-021-01134-x

40 Huang, W. et al. NFAT and NF-κB dynamically co-regulate TCR and CAR signaling responses in human T cells. Cell Rep 42, 112663 (2023). 10.1016/j.celrep.2023.112663

