## Supplemental Figures for "Multi-modal profiling identifies CD4+CXCR5+PD-1- Tfh cells as prognostic and predictive biomarkers for response to R-CHOP therapy in human DLBCL"

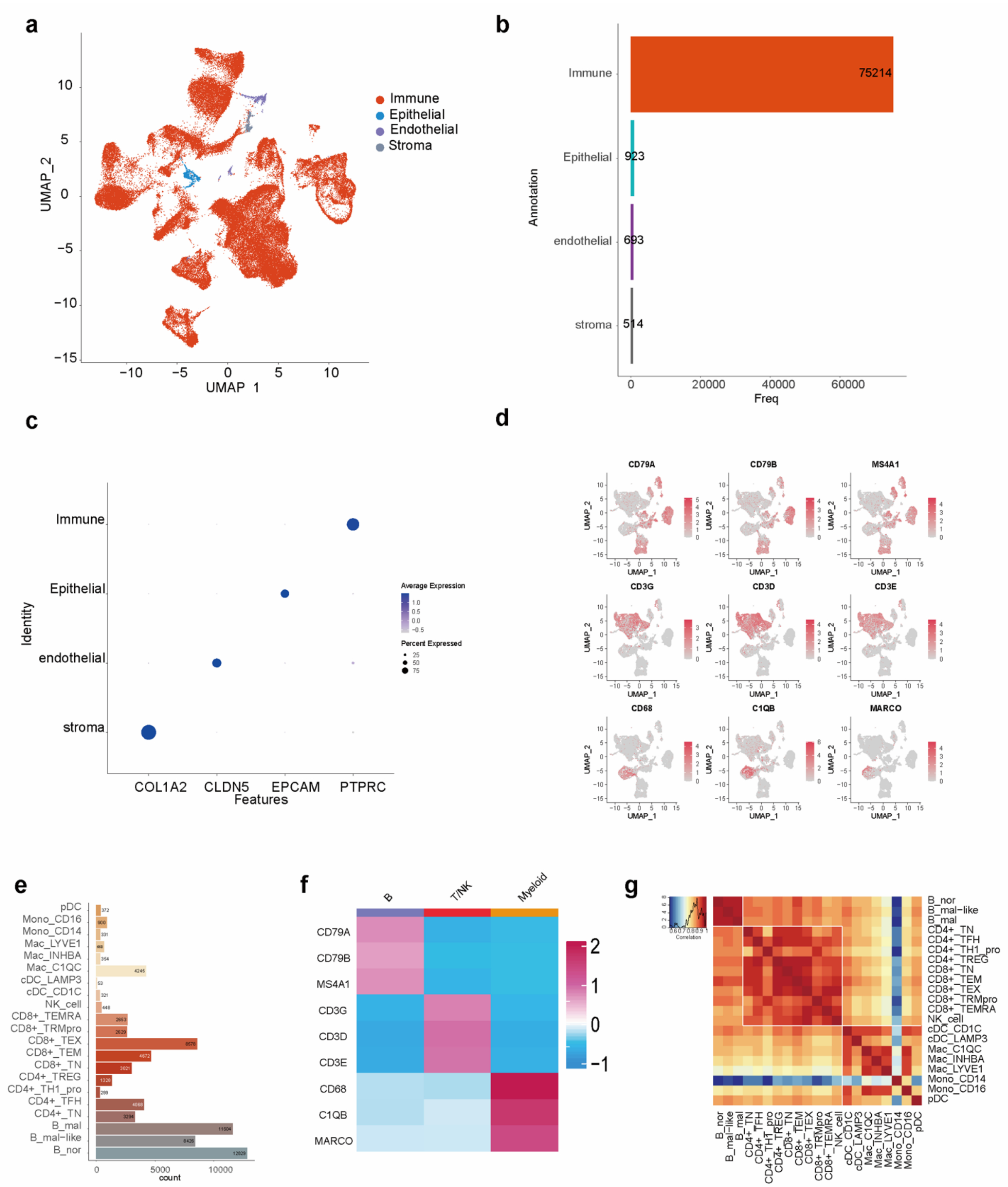


**Supplement Figure 1. Main classification for Single-cell profile of DLBCL.**

(a) UMAP plot of all cells colored by their cellular identity. (b) The bar plot showing the cell count of main cell types (epithelial cells [n = 923], immune cells [n = 75,212], stromal cells [n = 514], Endothelial [n=693]). (c) Canonical main cell type gene expression station is displayed by dot plot. (d) Canonical subpopulation cell type genes expression station is displayed by UMAP plot. (e) Bar plot showing the count of subpopulation cell types. (f) Heatmap showing the

Canonical subpopulation cell type genes expression. (g) Express correlation matrix confirmed the define of our work.

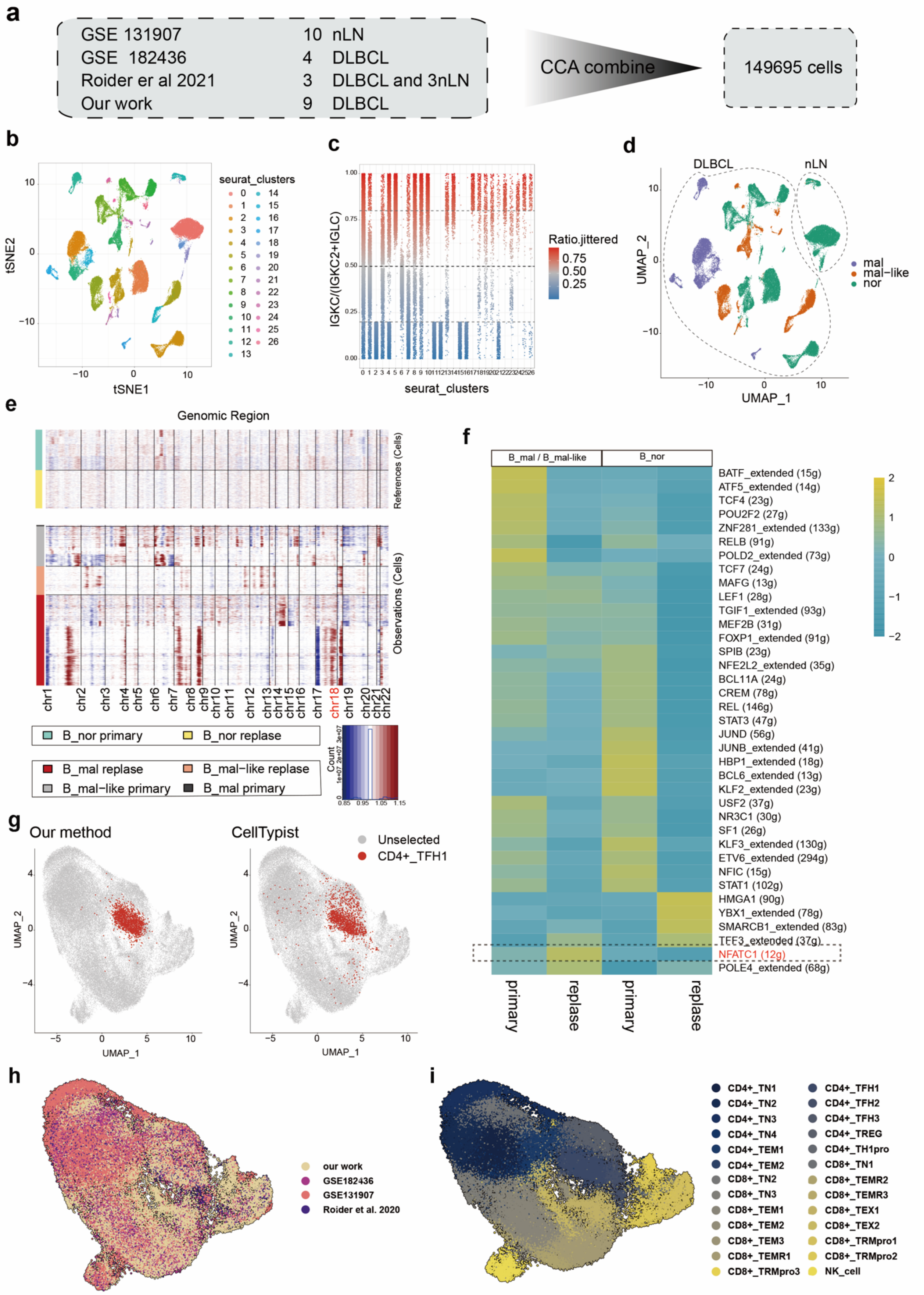

**Supplement Figure 2. Independent published Single-cell profile of DLBCL for validation our results.**

(a) This study applied canonical correlation analysis (CCA) method to combined our data and other independent DLBCL and nLN(normal lymph node) data. (b) UMAP demonstrates B cells in the combined single cell sequencing data, colored by graph-based clustering. (c) The scRNA expression profiles of the B cells were visualized by UMAP, colored by *IGKC* fraction. (d) B cells in the combined single cell sequencing data were visualized using UMAP, the cells are colored by new B cell classification. (e) Large-scale copy number alterations (CNAs) of B malignant or malignant-like and normal B cells identified using scRNA-seq data. Normal B cells were used as a reference. The CNA patterns for the B_mal and B_mal-like cell in the relapse or primary group, are shown. X axis, chromosome position; y axis, individual cells. Red color represents amplification group, while blue color represents deletion. (f) Coexpression modules in each cell cluster and their master regulators were identified using SCENIC. Color scale indicates regulon activity levels. (g) UMAP showing the comparation between our method (left) and CellTypist (right). T cells in the combined single cell sequencing data were visualized using UMAP, the cells are colored by data set names(h) and cell types (i).

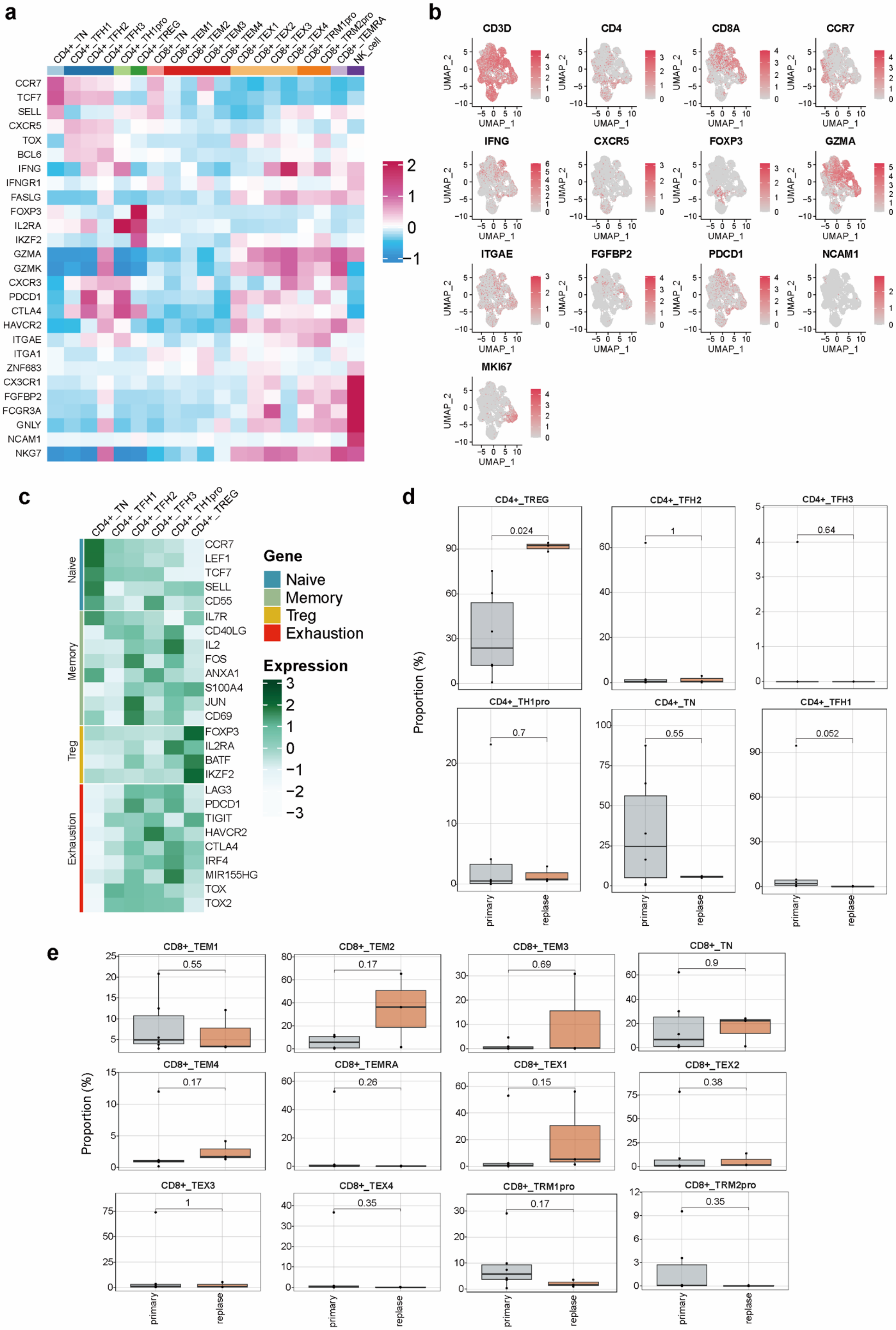
a
CD4+_TN
CD4+_TFH1
CD4+_TFH2
CD4+_TFH3
CD4+_TH1pro
CD4+_TREG
CD8+_TN
CD8+_TEM1
CD8+_TEM2
CD8+_TEM3
CD8+_TEM4
CD8+_TEX1
CD8+_TEX2
CD8+_TEX3
CD8+_TEX4
CD8+_TRM1pro
CD8+_TRM2pro
CD8+_TEMRA
NK_cell
CCR7
TCF7
SELL
CXCR5
TOX
BCL6
IFNG
IFNGR1
FASLG
FOXP3
IL2RA
IKZF2
GZMA
GZMK
CXCR3
PDCD1
CTLA4
HAVCR2
ITGAE
ITGA1
ZNF683
CX3CR1
FGFBP2
FCGR3A
GNLY
NCAM1
NKG7
b
CD3D
CD4
CD8A
CCR7
IFNG
CXCR5
FOXP3
GZMA
ITGAE
FGFBP2
PDCD1
NCAM1
MKI67
UMAP_1
UMAP_2
c
Naive
Memory
Treg
Exhaustion
CCR7
LEF1
TCF7
SELL
CD55
IL7R
CD40LG
IL2
FOS
ANXA1
S100A4
JUN
CD69
FOXP3
IL2RA
BATF
IKZF2
LAG3
PDCD1
TIGIT
HAVCR2
CTLA4
IRF4
MIR155HG
TOX
TOX2
Gene
Naive
Memory
Treg
Exhaustion
Expression
d
CD4+_TREG
0.024
CD4+_TFH2
1
CD4+_TFH3
0.64
CD4+_TH1pro
0.7
CD4+_TN
0.55
CD4+_TFH1
0.052
Proportion (%)
primary
replase
e
CD8+_TEM1
0.55
CD8+_TEM2
0.17
CD8+_TEM3
0.69
CD8+_TN
0.9
CD8+_TEM4
0.17
CD8+_TEMRA
0.26
CD8+_TEX1
0.15
CD8+_TEX2
0.38
CD8+_TEX3
1
CD8+_TEX4
0.35
CD8+_TRM1pro
0.17
CD8+_TRM2pro
0.35
Proportion (%)
primary
replase

7

**Supplement Figure 3. The classification of T cells in Single-cell profile of DLBCL.**

Canonical T cell subpopulation genes expression is displayed by heatmap (a) and UMAP (b). (c) The heatmap showing the functional gene expression on $CD4^+$ T cell cluster. (d) Relative contribution of each $CD4^+$ cell type (Percent of individual) is displayed by boxplot. (e) Relative contribution of each $CD8^+$ cell type (Percent of individual) is displayed by boxplot.

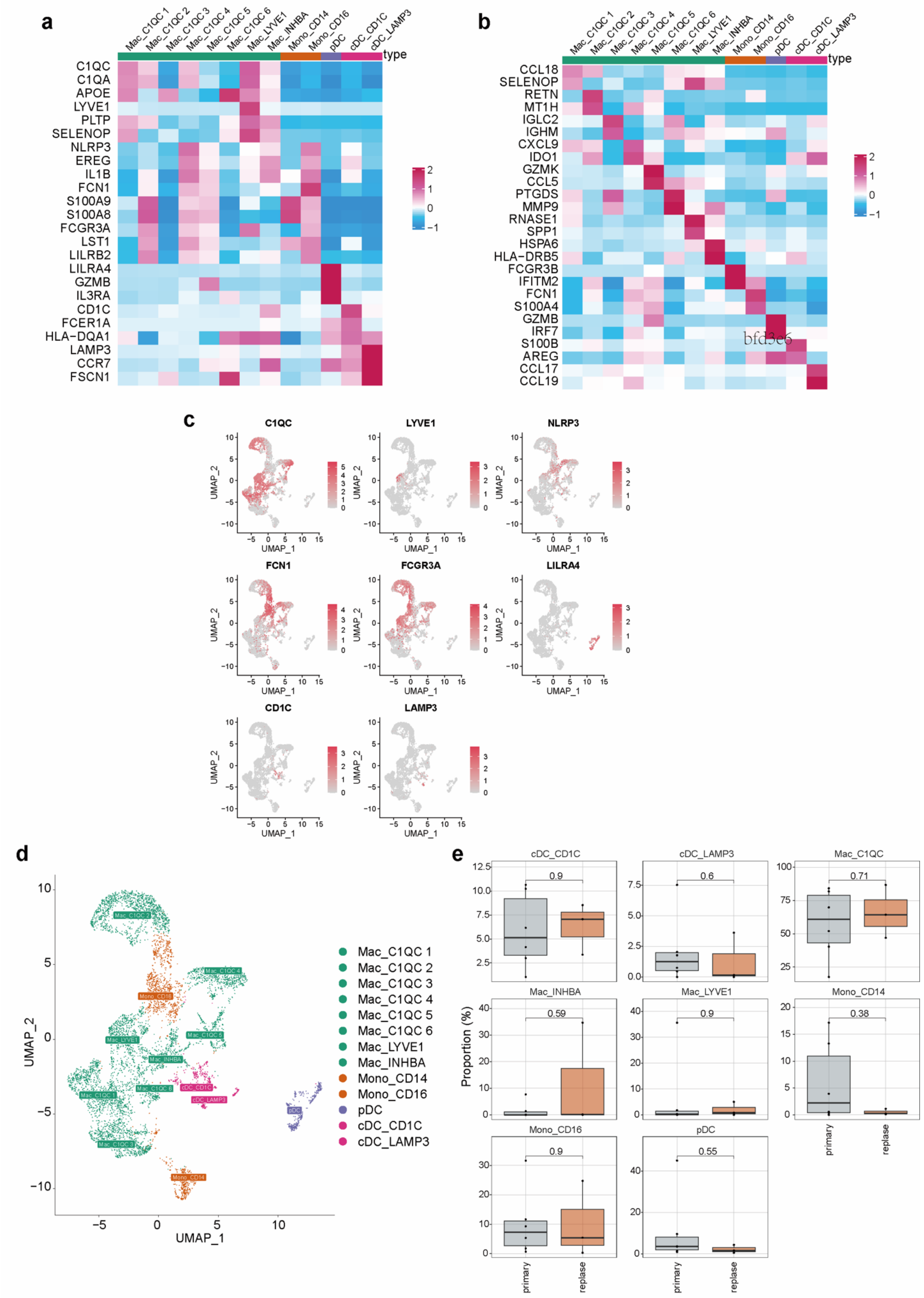


**Supplement Figure 4. The classification of Myeloid cells in Single-cell profile of DLBCL.**

(a) Canonical Myeloid cell subpopulation genes expression is displayed by heatmap and UMAP (b) Myeloid cell subpopulation novel specific genes are displayed by heatmap. (c) UMAP showing Canonical Myeloid cell subpopulation genes expression. (d) UMAP showing Myeloid cell subpopulation, colored by subpopulations names. (e) Relative contribution of each Myeloid cell subpopulation (Percent of individual) is displayed by boxplot.

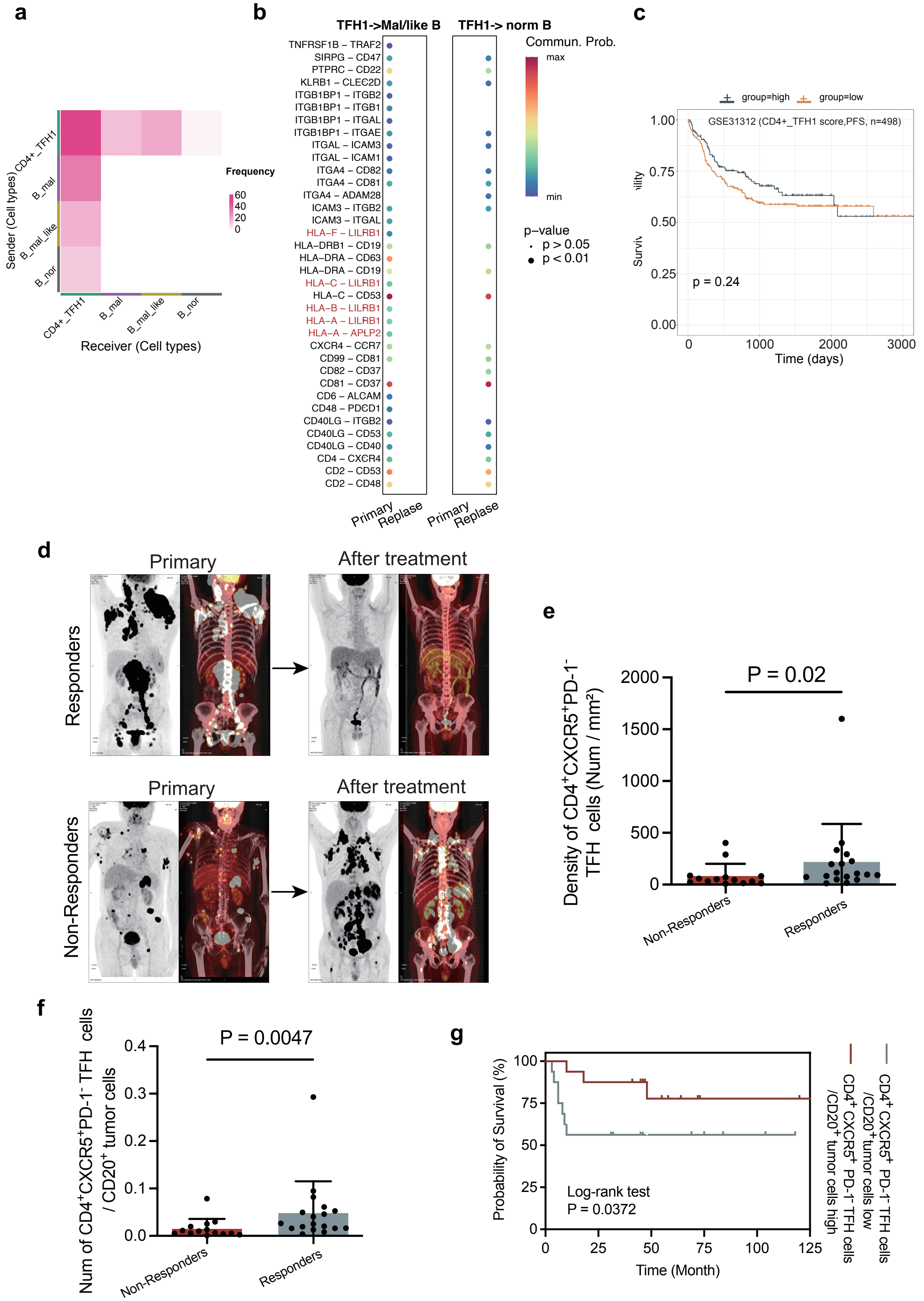


0
1 **Supplement Figure 5. Supplement Figure for figure 4 and figure 5.**

(a) The heatmap showing the all interaction pairs of $CD4^+$TFH1 and B cell types by LIANA with default database. (b)Dot plot showing the all interaction pairs of ligands of $CD4^+$TFH1 and receptors of B cell types by CellChat with cell-surface interactions in OmniPath. (c)Gene Set Variation Analysis (GSVA) to obtain $CD4^+$TFH1-specific gene scores. Subsequently, we divided the samples into two groups based on median and performed survival analysis using progression-free survival (PFS). (d) the representative PETCT images for responder or non- responder. (e) Density of $CD4^+CXCR5^+PD\text{-}1^-$ TFH cells, statistical significance was determined with the Wilcoxon signed-rank test. (f)Number of $CD4^+CXCR5^+PD\text{-}1^-$ TFH cells around every single $CD20^+$ tumor cells, statistical significance was determined with the Wilcoxon signed-rank test. (g) Overall survival analysis with number of $CD4^+CXCR5^+PD\text{-}1^-$ TFH cells around every single $CD20^+$ tumor cell in tissue samples from DLBCL patients.
